# Hispanic/Latino gastric adenocarcinoma patients have distinct molecular profiles including a high rate of germline *CDH1* mutations

**DOI:** 10.1101/764779

**Authors:** Sam C. Wang, Yunku Yeu, Suntrea T.G. Hammer, Shu Xiao, Min Zhu, Changjin Hong, Lynn Y. Yoon, Ibrahim Nassour, Jeanne Shen, Deepak Agarwal, Scott I. Reznik, John C. Mansour, Adam C. Yopp, Hao Zhu, Tae Hyun Hwang, Matthew R. Porembka

**Affiliations:** Department of Surgery, University of Texas Southwestern Medical Center, Dallas, Texas, USA; Children’s Research Institute, Departments of Pediatrics and Internal Medicine, Center for Regenerative Science and Medicine, University of Texas Southwestern Medical Center, Dallas, Texas, USA; Department of Quantitative Health Sciences, Lerner Research Institute, Cleveland Clinic, Cleveland, Ohio, USA; Department of Pathology, University of Texas Southwestern Medical Center, Dallas, Texas, USA; Department of Pathology, Stanford University School of Medicine, Stanford, California, USA; Department of Internal Medicine, University of Texas at Austin, Austin, Texas, USA; Department of Cardiovascular and Thoracic Surgery, University of Texas Southwestern Medical Center, Dallas, Texas, USA

**Author notes:** Equal contribution as primary authors (Sam C. Wang, Yunku Yeu). Equal contribution as senior authors (Tae Hyun Hwang, Matthew R. Porembka). **Corresponding Authors** 1. Sam C. Wang, Division of Surgical Oncology, Department of Surgery, The University of Texas Southwestern Medical Center, 5323 Harry Hines Blvd, Dallas, Texas, 75390-8548., 2. Tae Hyun Hwang, Department of Quantitative Health Sciences, Lerner Research Institute, Cleveland Clinic, 9500 Euclid Avenue Cleveland, Ohio 44195. **Author contributions** Study concept and design: SCW, YY, STGH, HZ, THH, MRP; acquisition of data: SCW, YY, STGH, SX, MZ, CH, LYY, IN, JS, DA, SIR, JCM, ACY, THH, MRP; analysis and interpretation of data: SCW, YY, STGH, CH, JS, HZ, THH, MRP; drafting of the manuscript: SCW, YY, THH, MRP; critical revision of the manuscript: all; obtained funding: SCW, HZ, MRP; study supervision: SCW, THH, MRP.

**Keywords:** Gastric cancer, Hispanic, Latino, cancer disparity, cancer genomics, *CDH1*, E-cadherin

## Abstract

Hispanic/Latino patients have a higher incidence of gastric cancer and worse cancer-related outcomes as compared to patients of other backgrounds. Whether there is a molecular basis for these disparities is unknown, as very few Hispanic/Latino patients were included in previous studies. We performed a large, integrated genomic analysis of gastric cancer samples from Hispanic/Latino patients. Whole-exome sequencing (WES) and RNA sequencing were performed on 57 Hispanic/Latino gastric cancer patient samples. Germline analysis was conducted on 83 patients. Functional testing of *CDH1* germline mutations was performed in Chinese hamster ovary cells. Tumors from Hispanic/Latino patients were significantly enriched for the genomically-stable subtype (as defined by The Cancer Genome Atlas), compared to Asians and Whites (65% vs 21% vs 20%, P < 0.001). Transcriptomic analysis identified molecular signatures that were prognostic. Of the 43 Hispanic/Latino patients with diffuse-type gastric cancer, 7 (16%) had germline mutations in *CDH1*. Mutation carriers were significantly younger than non-carriers (41 vs 50 years, P < 0.05). E-cadherin expression was reduced in 5 of 6 mutation carrier tumor samples available for analysis. *In silico* algorithms predicted 5 variants were deleterious. For the two variants that were predicted to be benign, we demonstrated that the mutations conferred increased migratory capability, suggesting pathogenicity. Hispanic/Latino gastric cancer patients possess unique genomic landscapes. This includes a high rate of *CDH1* germline mutations that may partially explain their aggressive clinical phenotypes. Individualized screening, genetic counseling, and treatment protocols based on patient ethnicity and race may be necessary.

## Introduction

Gastric cancer is the second-deadliest cancer worldwide, causing an estimated 834,000 deaths in 2016.^1^ Hispanic/Latino patients have different clinicopathologic features than patients of other ethnicities and races. In the United States, Hispanics/Latinos have twice the incidence and mortality from gastric cancer compared to non-Hispanic Whites.^2^ Hispanic/Latino gastric cancer patients also tend to be diagnosed at a younger age, with more advanced-stage disease, and with a higher proportion of diffuse-type cancers (DGC).^3–5^ While environmental exposures and socioeconomic factors likely contribute to the observed clinicopathologic differences, ethnicity/race-associated differences in tumor biology may also be involved. For example, African-American breast cancer patients have higher rates of triple-negative cancers and a higher prevalence of *TP53* mutations, as compared with White patients.^6, 7^

Whether there is a molecular basis for observed outcome differences for gastric cancer patients of different ethnicities/races has been heretofore unanswerable as previous large gastric cancer genomic studies had included very few Hispanic/Latino patients. The TCGA has performed the largest published sequencing study of gastric adenocarcinoma and included only five Hispanic/Latino patients in its 443-patient cohort.^8^ Other major sequencing efforts of gastric cancer originated in East Asia, including those by Ichikawa et al (207 patients) and Cristescu et al (300 patients); these studies also did not include any Hispanic/Latino patients.^9, 10^ Given the known association between ethnicity/race and tumor biology, the underrepresentation of Hispanic/Latino patients in previously published studies have likely biased our current genomic understanding of gastric cancer.^11^

To address this knowledge gap, we performed a large, integrated genomic analysis of samples from 83 Hispanic/Latino gastric cancer patients. Comparative analyses were performed using data from Asian and White patients previously published by The Cancer Genome Atlas (TCGA).^12^

## Methods

### Sample acquisition and processing

This study was approved by the University of Texas Southwestern Medical Center Institutional Review Board. All gastric adenocarcinoma patients who were self-reported as being of Hispanic/Latino ancestry were recruited to join the study. All enrolled patients provided written consent.

Blood samples were drawn and stored at −80°C prior to nucleic acid extraction. Tumor and adjacent non-neoplastic gastric tissue were obtained from treatment-naïve subjects via endoscopic biopsies or gastric resections. The samples were stabilized immediately in RNAlater (Ambion) for at least 24 hours at 4°C, then stored in liquid nitrogen until nucleic acid extraction. A second set of adjacent tissue samples from both the tumor and non-neoplastic stomach were also obtained for pathologic examination to confirm the histology, and to provide a microscopic assessment of tumor cellularity and extent of tumor necrosis. These samples were evaluated by a board-certified pathologist with expertise in gastrointestinal malignancies (S.T.G.H.) No samples were excluded on the basis of tumor cellularity, consistent with the TCGA collection protocol. Samples with greater than 10% necrosis were excluded. For some samples, RNA was isolated with mirVana miRNA Isolation Kits (Ambion) and DNA was isolated with QuickGene DNA Tissue Kits (Kurabo). Other samples were processed using the AllPrep DNA/RNA kits (Qiagen). Nucleic acid quality control was ensured with NanoDrop (Thermo Fisher) spectrophotometric quantitation and visualization on an agarose gel.

### Generation of *CDH1* mutants

Chinese hamster ovary (CHO) cells (ATCC, CCL-61) were maintained in F-12K medium (Gibco) with 10% fetal bovine serum supplementation. Wild-type human *CDH1* on a pcDNA3 plasmid was obtained (hE-cadherin-pcDNA3, Addgene, 45769). Variants were generated using the Q5 Site-Directed Mutagenesis Kit (New England Biolabs). Plasmid transfection was performed with Lipofectamine 3000 (Invitrogen). Selection was performed with G-418 (Sigma). Sanger sequencing was performed to confirm sequences using the following primers (Genewiz):

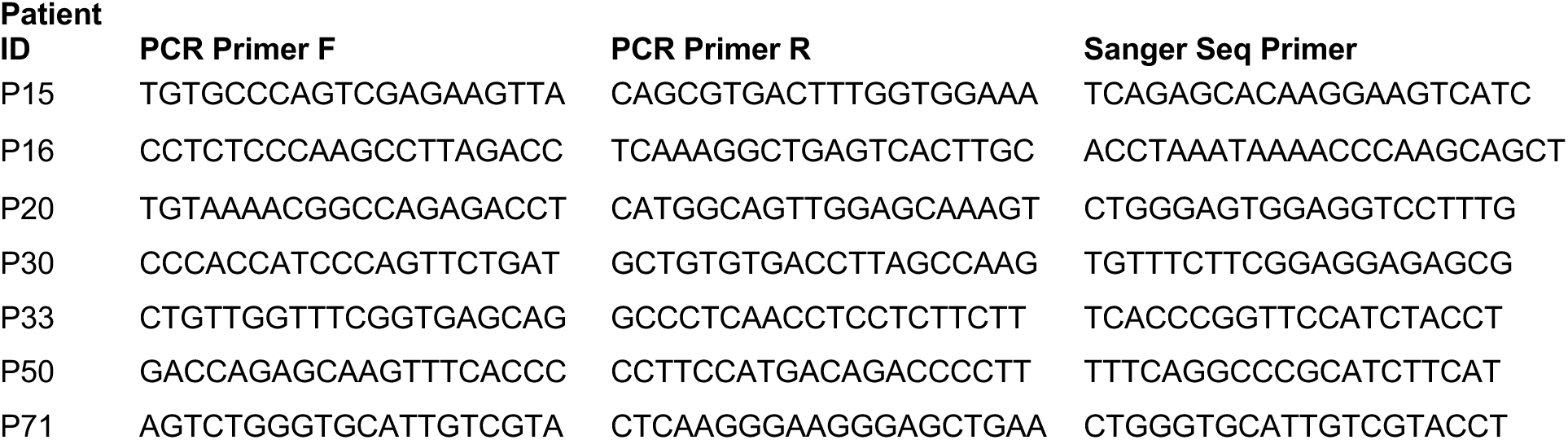

### Immunofluorescence

CHO cells were fixed on glass slides with pre-cooled methanol for 15 min at −20°C, and blocked by 1% bovine serum albumin in PBS-T for 1 hour at room temperature. The slides were then incubated with anti-E-cadherin antibody (Abcam, ab76055; 1:1000 dilution) at 4°C overnight, followed by secondary antibody at room temperature for 1 hour. DAPI was used as a nuclear stain (Vector Laboratories). Images were captured on a Zeiss confocal microscope.

### Immunohistochemistry

Antigen retrieval was performed with sodium citrate buffer, followed by incubation with anti-E-cadherin antibody (Abcam, ab76055; 1:1000) at 4°C overnight. Detection was performed with the ABC kit (Vector Laboratories) and DAB kit (Vector Laboratories) or MOM kit (Vector Laboratories).

### Scratch assay

CHO cells transfected with a given plasmid were grown to confluence. A scratch was made and three images were taken of each well. 24 hours later, three more images of each well were taken. The distance between the wound edges was measured using cellSens Dimension software (Olympus). The average of the three images from each time point was used as one biological replicate. Two independent experiments with at least four biological replicates for each genotype were performed.

### Statistical analysis

The Mann-Whitney U test was used to compare continuous variables. Categorical variables were presented as counts and proportions and compared with Fisher exact tests. Survival was estimated using the Kaplan-Meier method and compared via the log-rank test. For the epidemiologic studies, data was presented as medians with interquartile ranges and full ranges in box and whisker plots and compared with the Kruskal-Wallis test.

### Whole-exome sequencing, RNA sequencing, and bioinformatic analyses

Please see the Supplementary Methods section for details regarding the whole-exome sequencing (WES), RNA sequencing (RNA-seq), and bioinformatic analyses. All sequencing data will be deposited into the Gene Expression Omnibus (GEO) upon acceptance or as needed for review.

### Patient and public involvement statement

Neither patients nor the public were involved in the design, conduct, reporting, or dissemination of our research.

## Results

We performed WES and RNA-seq on tissue samples from 57 patients. Blood samples were also obtained from 52 of these patients and used as normal controls. For the five patients for whom blood samples were unavailable, we used non-neoplastic gastric tissue as controls. We also performed WES on blood samples from an additional 26 patients (Table 1 and Supplementary Table 1). The mean coverage for WES was 267x for the 57 tumor samples, 209x for the five non-neoplastic gastric samples, and 67x for the 78 blood samples. For RNA-seq, the average number of reads was 96.9 million, with an average mapping rate of 97.6% for the 57 tumors and 5 non-neoplastic gastric samples.

**Table 1:**
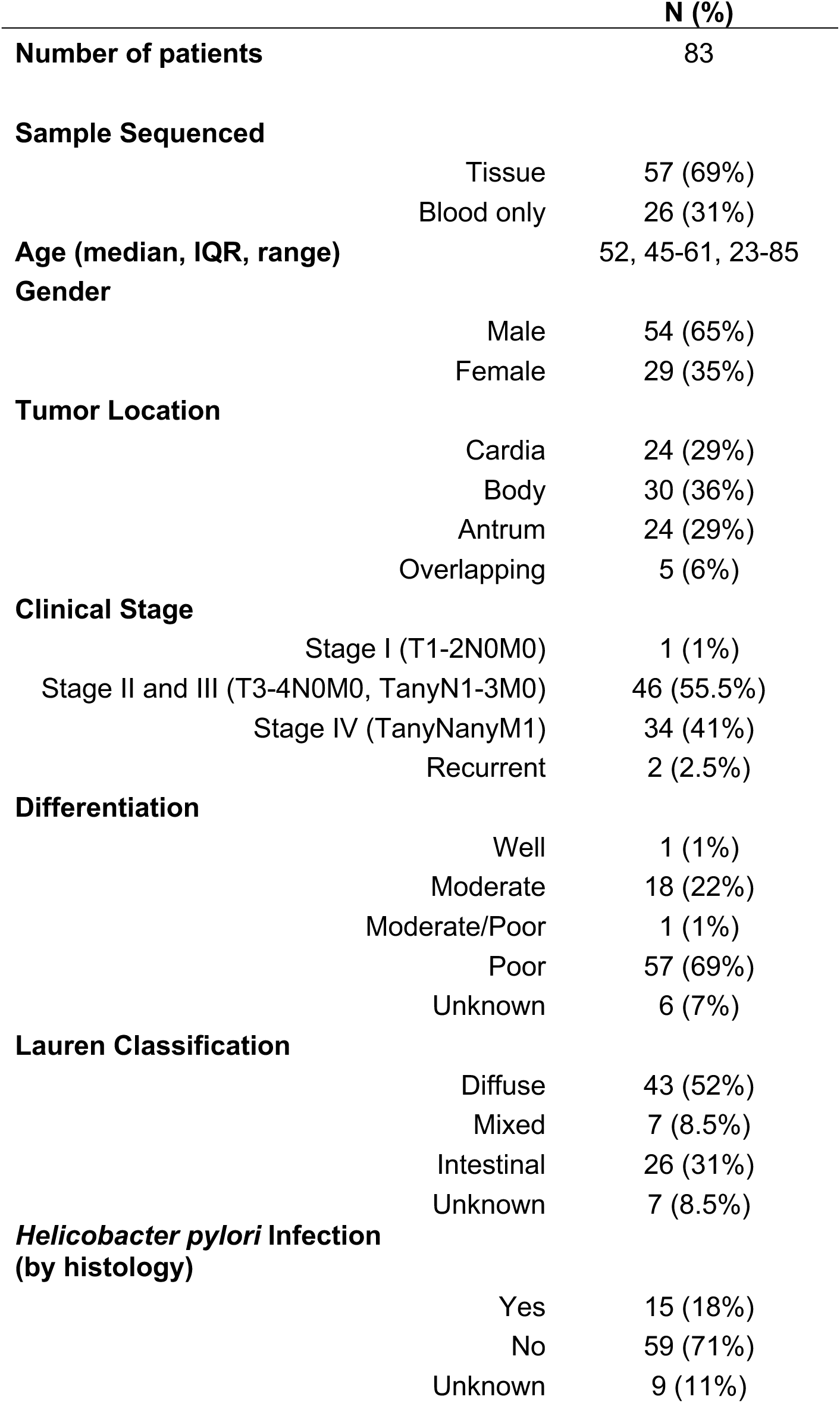
Clinicopathologic characteristics of Hispanic/Latino gastric cancer patients in this study. IQR: interquartile range.

Consistent with previous reports, the median age at time of diagnosis for the 83-patient Hispanic/Latino patient cohort was younger than that for the 77 Asian and 172 White patients analyzed by the TCGA (52 years, vs 66 and 66 years, respectively, *P* < 0.0001, Supplementary Fig. 1a). To confirm that the Hispanic/Latino cohort’s self-reported ancestry was unique from that of the TCGA Asian and White patients, we compared WES data from each of the three groups to reference data available through the Human Genome Diversity Project (HGDP).^13^ Using principal component analysis, we found that the Hispanic/Latino cohort clustered independently from the Asian and White patients in the TCGA groups and were related most closely to the HGDP samples from Central and South America (Fig. 1 and Supplementary Fig. 1b).

**Figure 1.**
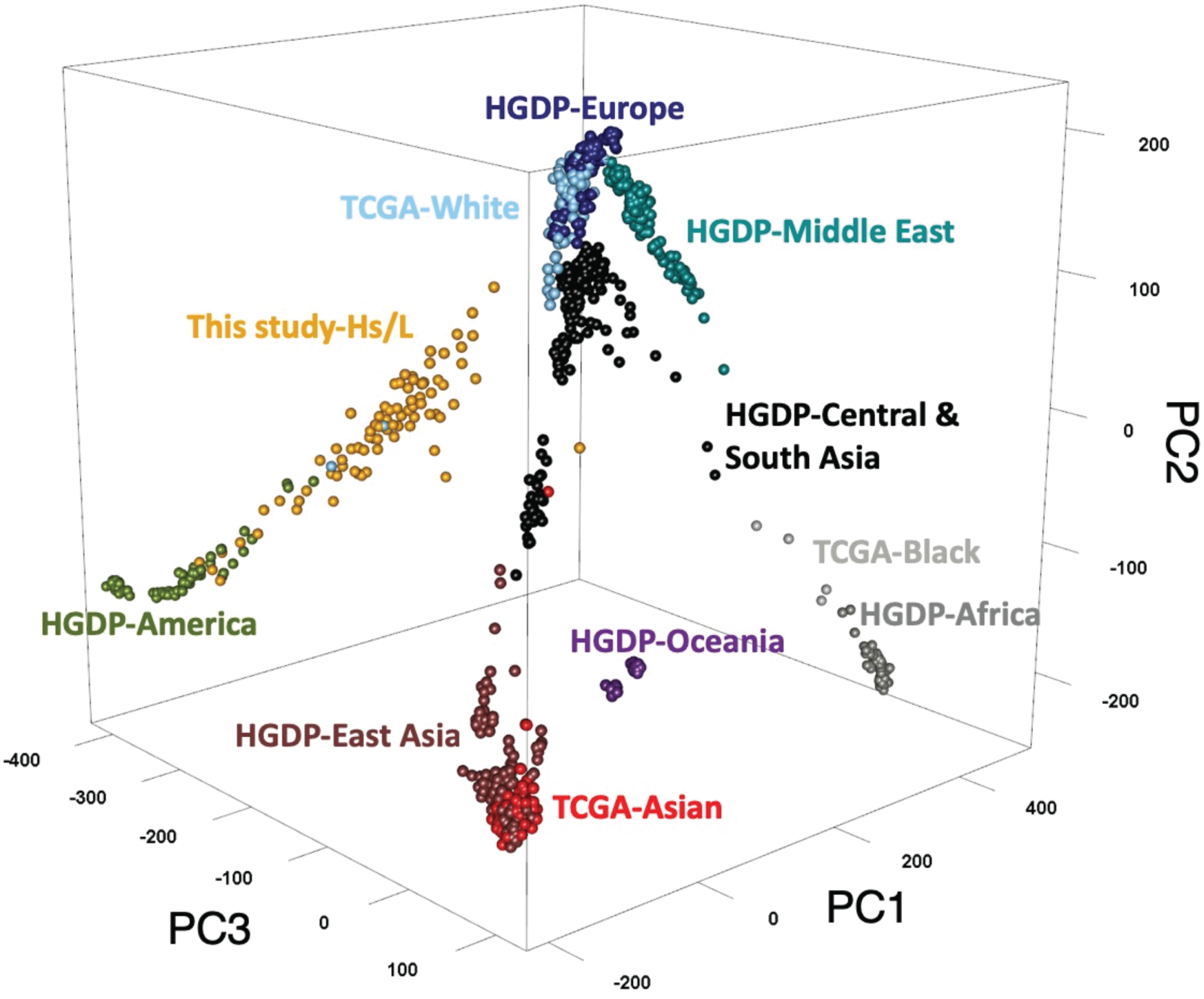
Hispanic/Latino (Hs/L) gastric cancer patients are of unique ancestry as compared to Asian and White patients. Principal component (PC) analysis was performed using Locating Ancestry from SEquence Reads (LASER) comparing Hispanic/Latino patients from this study to Asian and White patients analyzed by The Cancer Genome Atlas (TCGA). The Human Genome Diversity Project (HGDP) was used as the reference.

### Gastric Cancers in Hispanic/Latino Patients are Enriched for the Genomically-Stable Subtype

We next classified the 57 Hispanic/Latino gastric cancer samples into one of the four molecular subtypes established by the TCGA (Supplementary Fig. 2a).^12^ We did not include African-Americans in this analysis, as there were only four African-American patients in the TCGA cohort. Tumors were first characterized based on Epstein-Barr virus (EBV) infection status, which was determined bioinformatically with PathoScope 2.0.^14^ We found no EBV infections, whereas 10% of the TCGA cohort was EBV-positive.^12^ Next, microsatellite instability (MSI) was assessed bioinformatically using MSISensor, which has previously demonstrated near-perfect concordance with the results of PCR or immunohistochemical analysis.^15, 16^ Three of the 57 samples (5%) had MSIsensor scores of greater than 10, indicating microsatellite instability (Supplementary Fig. 2b). Accordingly, these three samples showed mutation burdens greater than 13 mutations per megabase (Mb), whereas the average mutation burden for the 54 non-MSI samples was 2.5 mutations per Mb.

The remaining samples underwent somatic copy number alteration (SCNA) analysis.^10^ 17 samples (30%) had high SCNA scores, which placed them into the CIN group, and 37 patients (65%) had low scores and were categorized as genomically stable (GS; Fig. 2a). When compared to the Asian (20%) and White (21%) patients, Hispanic/Latinos had a significantly higher proportion of GS tumors (65%, P < 0.001; Fig. 2b). There were no significant differences between Asian and White patients in the proportions of subtypes. CIN samples showed an average of 3.5 mutations per Mb, while GS tumors had 2.0 mutations per Mb. This is consistent with the TCGA data found on the Broad Firehose, which showed CIN and GS samples as having 3.3 and 1.8 mutations per Mb, respectively (http://firebrowse.org/?cohort=STAD).

**Figure 2.**
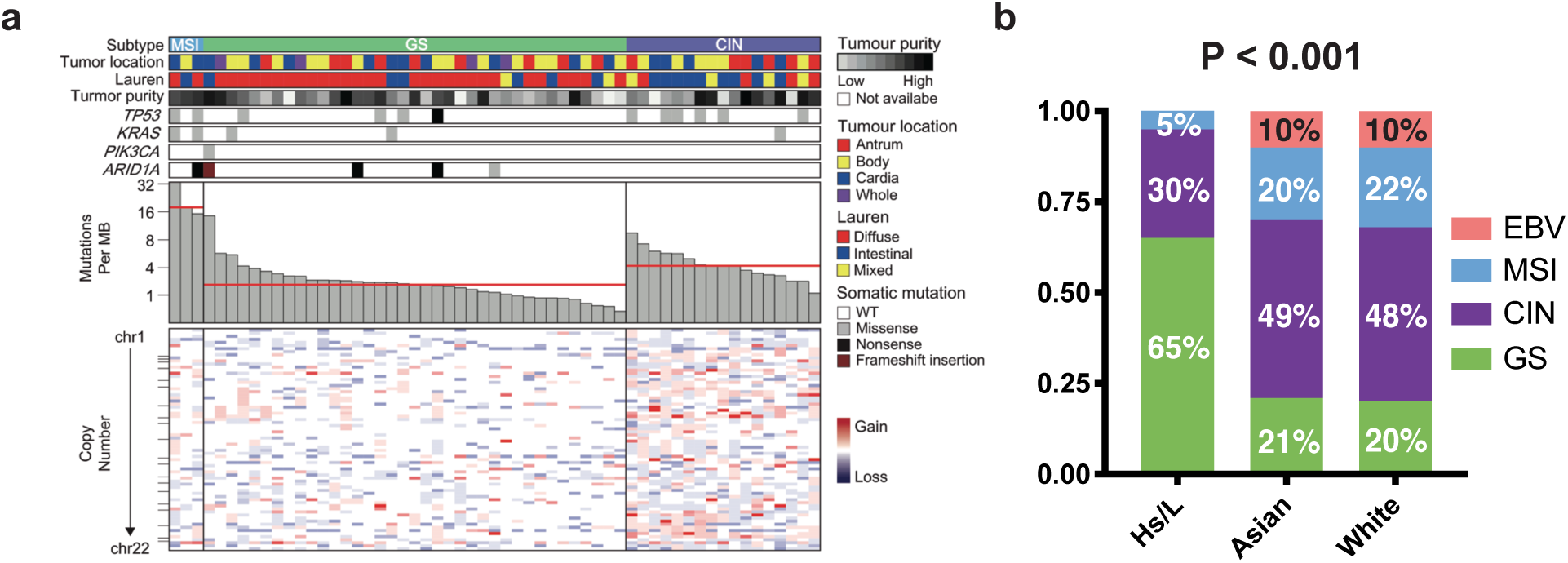
Gastric cancer in Hispanic/Latino (Hs/L) patients are predominantly of the genomically stable subtype. a. Tumors from 57 Hispanic/Latino gastric cancer patients were subtyped and listed by descending mutation burden. Clinical and molecular data are depicted. MB = megabase, WT = wild-type. b. Molecular classification of samples within each ethnicity/race. P < 0.001. EBV: Epstein-Barr virus infected, MSI: microsatellite instability, CIN: chromosomal instability, GS: genomically stable.

In the TCGA analysis, the GS subtype was found to be enriched for tumors with diffuse-type histology.^12^ Accordingly, we found that of the 37 GS patients, 78% had diffuse-type, 16% had intestinal-type, and 6% had mixed-type tumors. In contrast, the CIN cohort was comprised of 23.5% diffuse, 53% intestinal, and 23.5% mixed-type tumors (*P* < 0.001, Fig. 2a).

### Hispanic/Latino Gastric Cancers Recapitulate Key Genomic Features Identified by the TCGA

Although the Hispanic/Latino samples were significantly enriched for GS tumors, many defining genomic alterations previously identified by the TCGA were recapitulated in the current cohort. For example, the most common recurrent mutation in Hispanic/Latino gastric cancer samples was *TP53*. Other driver mutations commonly associated with gastric cancer, such as those in *ARID1A* and *APC*, were also found at a comparable frequencies (Fig 3a). We also identified similar structural variations. The TCGA identified *CLDN18-ARHGAP* fusions in 15% of their GS-type tumors. These rearrangements lead to dysregulated *RHOA* signaling and loss of an epithelial phenotype.^12, 17^ Using FusionCatcher and STAR-fusion to evaluate our RNA-seq data, we found that four (11%) tumors had this rearrangement (Fig. 3b and Supplementary Table 2).^18, 19^ All four were GS and diffuse-type. We also observed that 76% of CIN tumors, which were enriched for intestinal-type histology, had amplifications in the 8q24.21 region. This was significantly higher than seen in GS samples, in which only 19% had this copy number abnormality (Fig. 3b; *P* < 0.001). We confirmed this finding in the TCGA cohort, which similarly showed an enrichment of this structural alteration in CIN samples.^12^ The 8q24.21 region most notably carries the *MYC* oncogene, and other groups have noted that amplifications in this region are common in intestinal-type gastric cancers and are associated with worse outcomes in gastric cancer patients.^20–22^ Finally, we identified five instances of *KRAS* amplification (12p12.1), all of which occurred in CIN patients, consistent with previous reports (Fig. 3b).^23^

**Figure 3.**
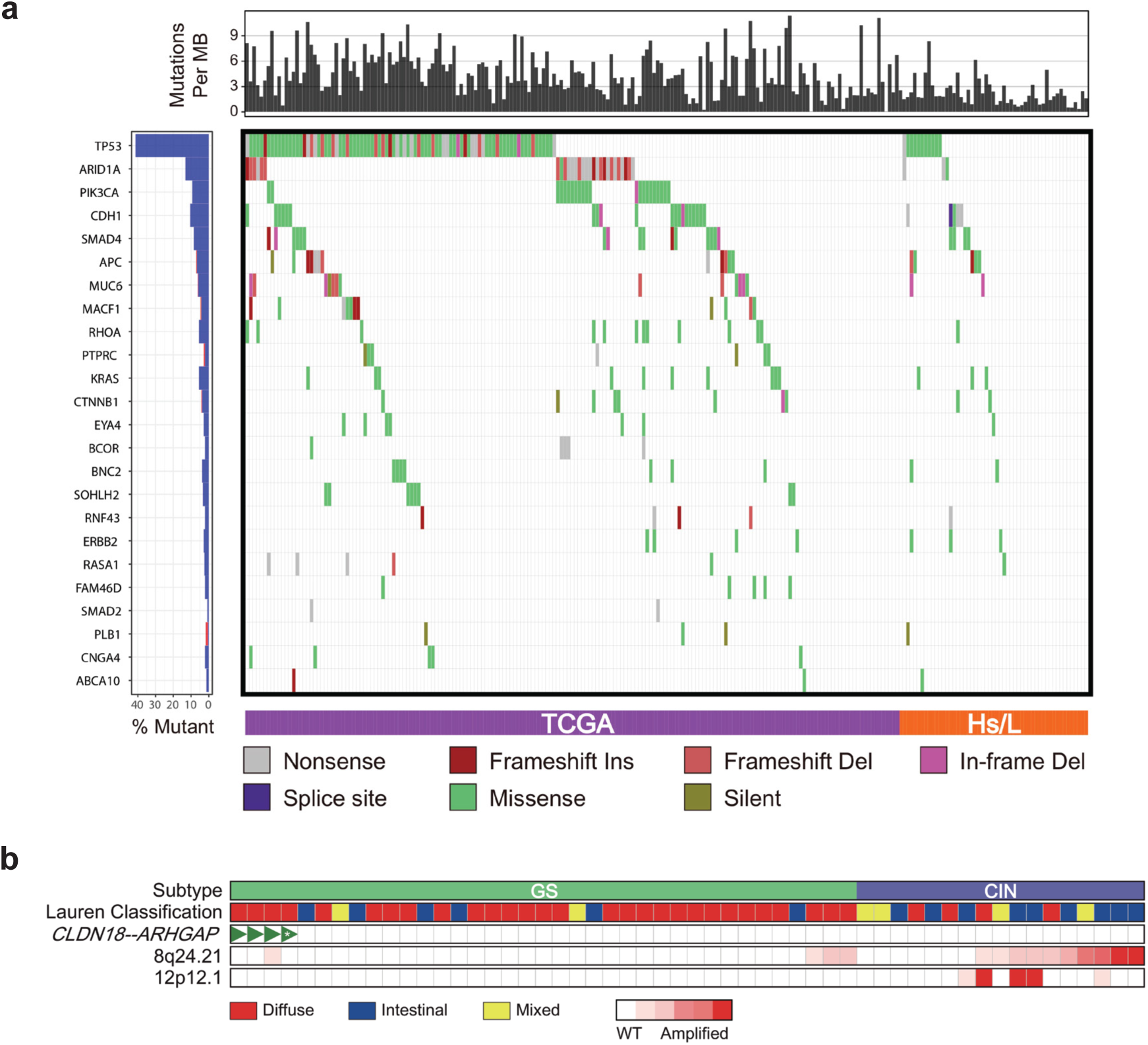
Key genomic features of gastric cancer are identified in Hispanic/Latino (Hs/L) samples. a. Recurrent somatic mutations identified by the TCGA in non-hypermutated gastric cancer samples from Hispanic/Latino patients. b. Structural variations seen in Hispanic/Latino gastric cancer samples. CIN: chromosomal instability, GS: genomically stable, WT: wild-type, *: CLDN18-ARHGAP45, all other fusions were CLDN18-ARHGAP26.

### Gene Expression Profiling is Prognostic

To further interrogate the RNA-seq dataset, we selected top 50 most variably expressed genes and performed consensus clustering to identify patient subgroups. We found five clusters with distinct clinicopathologic profiles (Fig. 4a, Supplementary Table 3). Patients in Clusters 2 and 3 tended to be younger, while Clusters 1 and 5 patients were older. Cluster 3 patients had tumors enriched for diffuse-type and GS tumors. We also found that the clustering provided significant prognostic capability. Cluster 1 patients had the shortest median survival at 7.7 months, whereas Cluster 4 patients had the longest survival, with median survival not reached. Patients in Clusters 2, 3, and 5 had intermediate risk profiles with median survival of 19.7 months (Figure 4b, P < 0.001; Supplementary Fig. 3a, P < 0.01). Importantly, the prognostic value of mRNA clustering was maintained when patients were stratified by molecular subtype or by Lauren classification (Supplementary Fig 3b-d, P < 0.05 for each). When we performed Gene Set Enrichment Analysis^24^ to identify pathways that were uniquely overexpressed in Cluster 1 and 4 tumors, we found that the upregulated pathways in Cluster 1 were involved in cell cycle regulation, cell growth, and epithelial-mesenchymal transition (Fig. 4c) while upregulated pathways in Cluster 4 were associated with an activated immune response (Fig. 4d).

**Figure 4.**
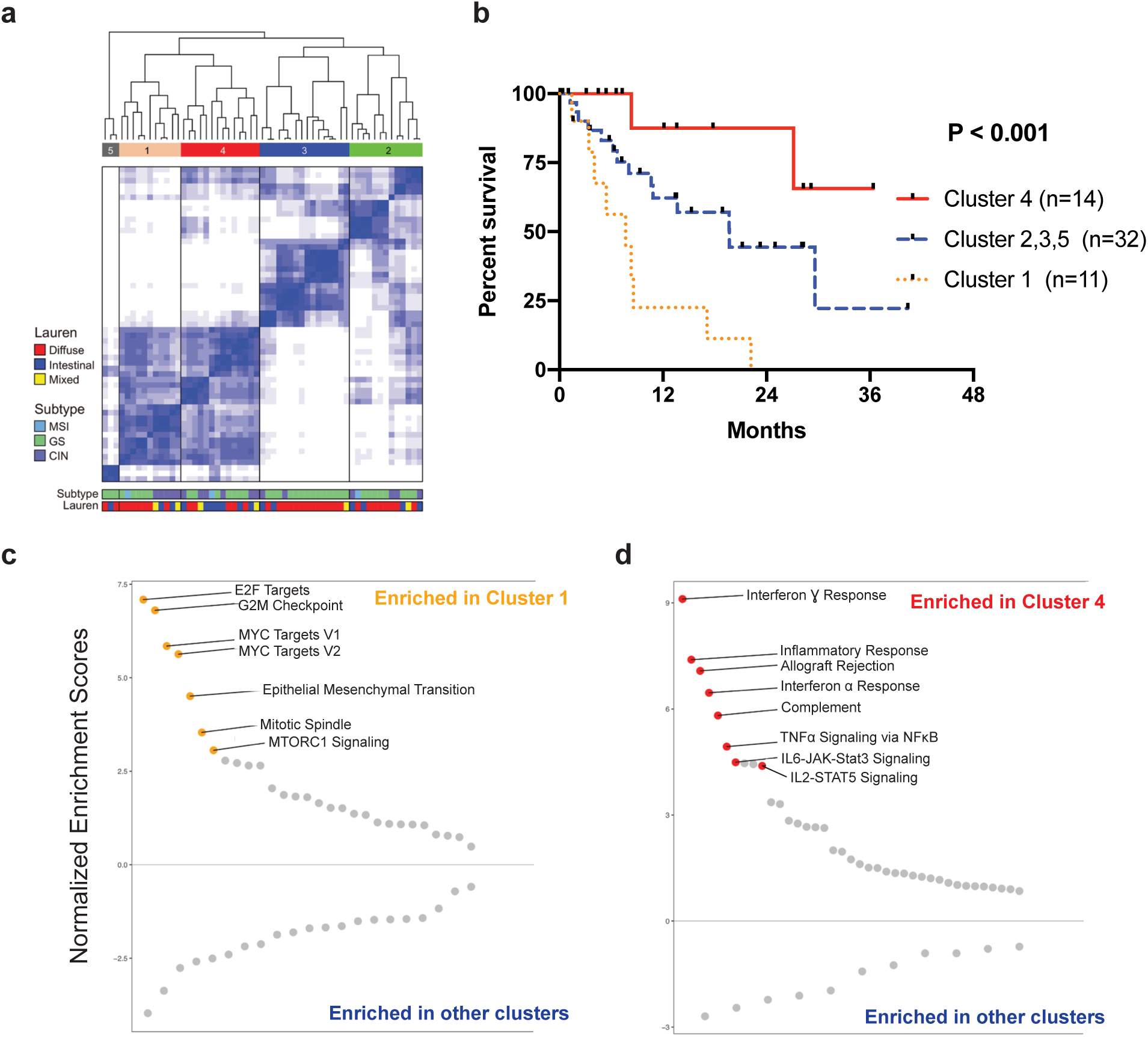
Transcriptomic signatures of gastric cancer from Hispanic/Latino patients are prognostic. a. Unsupervised consensus clustering based on the top 50 most variably expressed genes. MSI: microsatellite instability, CIN: chromosomal instability, GS: genomically stable. b. Kaplan-Meier curve comparing overall survival based on clusters. P < 0.001. c. Normalized enrichment scores from Gene Set Enrichment Analysis (GSEA) comparing Cluster 1 to Clusters 2, 3, 4, and 5. Orange dots denote Hallmark gene sets related to cell cycle, cell growth, and epithelial-mesenchymal transition, all of which had false-discovery rate q-value < 0.01 d. Normalized enrichment scores from GSEA analysis comparing Cluster 4 to Clusters 1, 2, 3, and 5. Red dots denote immune-related Hallmark gene sets, all of which had false-discovery rate q-value < 0.01.

### Hispanic/Latino Patients with Diffuse-Type Tumors Have Frequent Germline CDH1 Mutations

We analyzed the WES data from either blood or non-neoplastic stomach from 83 patients and identified seven germline *CDH1* mutations (Fig. 5a, Table 2). All seven mutations were identified in patients with diffuse-type cancer (DGC; 16%) and were confirmed with Sanger sequencing (Supplementary Fig. 4, Supplementary Table 4). Two mutations were deletions and five were missense variants. In patients with DGC, the median age of mutation carriers was 41 years (range 36-54 years) while the median age of *CDH1* wild-type patients was 50 years (range: 26-76 years; P < 0.05; Supplementary Table 1).

**Figure 5.**
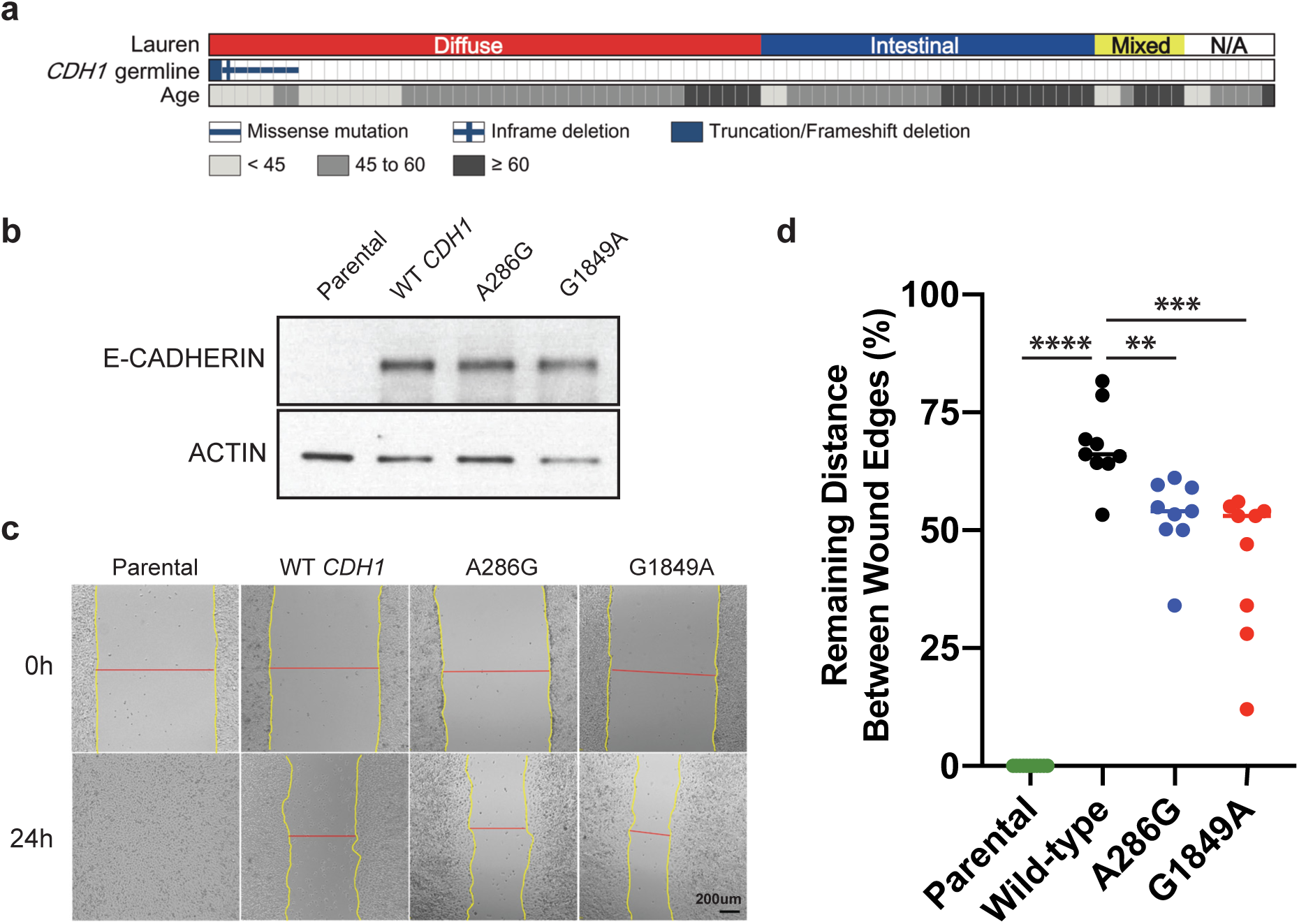
Hispanic/Latino gastric cancer patients have high rates of germline *CDH1* mutations. a. Seven germline *CDH1* mutations were identified in patients with diffuse gastric cancer. b. Western blot showing E-cadherin expression level upon transfection of plasmids carrying wild-type *CDH1*, A286G variant, or G1849A variant into Chinese hamster ovary (CHO) cells. c. Representative pictures of scratch assays. Distance between the wound edges were measured after 24 hours. d. Quantification of remaining distance between wound edges, relative to 0h. N ≥ 9 per group, with at least two independent experiments. ** P < 0.01, *** P < 0.001, **** P < 0.0001

**Table 2:** Germline *CDH1* mutations found in the Hispanic/Latino gastric cancer cohort.

| Age | Lauren | Chromosome position | Amino Acid Changes | Mutation Type | ClinVar | Predicted Effect by SIFT & PolyPhen-2 | gnomAD | gnomAD (Latino) |
| --- | --- | --- | --- | --- | --- | --- | --- | --- |
| 51 | Diffuse | Exon3:c.286A>G | I96V | missense | Benign | Benign | 0.016% | 0.12% |
| 36 | Diffuse | Exon13:c.1988_2011del, 2012A>C | 663-671del<br>EVGDYKINLKLMND<br>QN>VGDSNQND | deletion - inframe and missense | N/A | Deleterious | . | . |
| 37 | Diffuse | Exon12:c.1849G>A | A617T | missense | Benign | Benign | 0.45% | 0.30% |
| 41 | Diffuse | Exon14:c.2276G>C | G759A | missense | Uncertain | Deleterious | 0.0004% | 0.0029% |
| 40 | Diffuse | Exon2:c.135delC | H45fs | deletion - frameshift | N/A | N/A | . | . |
| 41 | Diffuse | Exon6:c.715G>A | G239R | missense | Conflicting | Deleterious | . | . |
| 54 | Diffuse | Exon16:c.2558C>T | S853L | missense | Uncertain | Deleterious | 0.0016% | 0.009% |

Pathogenic *CDH1* germline mutations are known to cause hereditary DGC syndrome. However, none of the germline mutation carriers in our Hispanic/Latino cohort had a family history of gastric cancer or lobular breast cancer, which is another manifestation of the mutation.^25^ Previous reports have suggested that germline *CDH1* mutations contribute to early-onset gastric cancer in patients without family histories of cancer.^26^ We performed a literature search to estimate the rate of germline *CDH1* mutations in gastric cancer patients without family histories of gastric cancer, and identified four studies with relatively large cohorts. These included patients from Italy,^27^ Canada,^28^ China,^29^ and Korea.^30^ Out of a total of 600 patients, 350 had DGC, and approximately 12 germline mutations in the coding region of *CDH1* were identified across 13 patients (3.7%), with 3 patients having deletions, and 10 having missense alterations (Supplementary Table 4).^31, 32^ Thus, the prevalence of germline *CDH1* mutations in patients without a relevant family history was markedly higher in the Hispanic/Latino cohort than what has been reported in other ethnic/racial groups.

To determine whether the identified *CDH1* mutations were pathogenic, we first checked the population frequency of these variants in the Genome Aggregation Database (https://gnomad.broadinstitute.org). All seven variants were found in less than 1% of both the general population and in the Latino cohort (Table 2). We next queried the annotations of the five missense mutations in the ClinVar database.^33^ Two were classified as benign (P15 and P20) while the rest were either of uncertain significance or had conflicting data (P30, P50 and P71). Next, we used SIFT and PolyPhen-2 to predict the variant functionality via a bioinformatic approach.^31, 32^ Consistent with ClinVar annotation, P15 and P20 were predicted to be benign, but P30, P50, and P71 were projected to be pathogenic (Table 2). Of the six patients whose tissue samples were available for analysis, we performed immunohistochemistry for E-cadherin, and found that there was either decreased or complete loss of protein expression in five of the six patients, including in P15 and P20, who harbored putatively benign variants (Supplementary Fig. 5).

The variant found in P15, who was a 51 year-old man presenting with locally advanced disease, was a c.286 A>G transition that resulted in an I96V amino acid alteration. Patient 20, who was a 37 year-old woman presenting with metastatic disease, had an c.1849 G>A change that led to an A617T amino acid change. To test the effects of these variants *in vitro*, we generated plasmids carrying wild-type *CDH1* or these two variants and transfected them into Chinese hamster ovary (CHO) cells, which do not express E-cadherin at baseline and has been used extensively by other groups to test *CDH1* variant function^34–36^ Sanger sequencing confirmed that the mutations were generated correctly. We found that both 286 A>G and 1849 G>A variants generated protein products that were normal in size and cellular localization (Fig. 5b and Supplementary Fig. 6).

E-cadherin is involved in cell-cell adhesion and its loss can result in increased cellular migration. We performed scratch assays to test if 286 A>G or 1849 G>A affected the migratory ability of CHO cells. After 24 hours, parental CHO cells had completely covered the scratch. As expected, CHO cells expressing wild-type *CDH1* led to significantly reduced cellular migration, with 68% of the wound distance remaining (P < 0.0001). However, 286 A>G expressing cells had only 54% (P < 0.01) and 1849 G>A expressing cells had only 53% (P < 0.001) of their wound distances remaining (Fig. 5c and 5d). Thus, both variants conferred significantly increased migratory capability.

## Discussion

Hispanic/Latino patients experience significant gastric cancer outcome disparities. Whether there is a molecular basis for these differences is unknown, as previous gastric cancer genomic studies included very few Hispanic/Latino patients.^9, 10, 12^ To our knowledge, the only study to date that had included a large number of Hispanic/Latino patients was performed by Sahasrabudhe et al.^37^ However, their analysis of 333 patients from Latin America was limited to targeted sequencing of only five genes involved in DNA repair.

In this study, we have performed a large, integrated analysis of Hispanic/Latino gastric cancer samples and compared our results to those from Asian and White patients samples published by the TCGA. We found that Hispanic/Latino gastric cancer patients had a high incidence of germline *CDH1* mutations, and that their tumors were enriched for the GS molecular subtype. Our findings suggest that the lack of ethnic and racial diversity in samples analyzed by previous large-scale studies has likely biased our genomic understanding of gastric adenocarcinoma due to the overrepresentation of White and Asian patients. Previous studies in other cancer types also identified genomic differences based on ethnicity and race. Shi et al found that more than 50% of Asian non-small-cell lung adenocarcinoma patients had *EGFR* mutations, as compared to 20% of White patients.^39^ In a study of African-American prostate cancer patients, Yamoah et al identified genomic biomarkers related to race that were highly prognostic.^40^ Thus, having ethnically and racially representative study cohorts will enhance our understanding of fundamental disease biology and ensure that the efficacy of a selected treatment has been tested and confirmed for the patient’s ethnic/racial background.^11, 38^ Improved recruitment of underrepresented patient populations into future clinical and basic scientific studies should be mandatory.

The high rate of germline *CDH1* mutations in our Hispanic/Latino DGC cohort is striking. Of the seven mutations we identified, which represented 16% of the DGC patients, two had not been previously reported in ClinVar, three were annotated as uncertain or conflicting, and two were designated as benign. Thus, these variants would likely have been excluded as pathogenic. However, several lines of evidence suggest that these mutations have deleterious effects. First, previous studies have suggested that germline *CDH1* mutations may contribute to early-onset DGC.^26^ The variant carriers in our cohort had a median age of diagnosis of 41 years as compared to DGC patients with wild-type *CDH1* who had a median age of 50 at diagnosis. Second, E-cadherin protein expression was abnormal in five of the six tumors from *CDH1* mutation carriers that were available for analysis. Third, *in silico* analysis predicted that three of the five missense mutations were pathogenic. Finally, functional modeling of the two missense variants annotated by ClinVar and predicted to be benign by both SIFT and PolyPhen2 demonstrated pathogenic cellular migration phenotype. Our findings speak to the limitations of the currently available tools to predict accurately the pathogenicity of a given variant. When germline *CDH1* mutations are identified in patients who have a high pre-test probability of carrying a pathogenic variant, such as in a young DGC patient, more rigorous functional testing should be utilized to determine pathogenicity.

Germline *CDH1* mutations are one of the causes hereditary DGC syndrome. Since none of the seven Hispanic/Latino *CDH1* variant carriers had a family history of gastric cancer or lobular breast cancer, these mutations are either *de novo* or exhibited low penetrance. Previous estimates suggesting that carriers of pathogenic *CDH1* variants have a lifetime risk of up to 80% of developing DGC are likely overestimations as they are based on families that fulfil the International Gastric Cancer Linkage Consortium (IGCLC) guidelines and thus subjected to ascertainment bias.^41^ Recent studies that examined DGC penetrance in carriers of pathogenic *CDH1* variants that do not fulfil IGCLC criteria suggest a lower lifetime gastric cancer risk. Xicola et al found a lifetime risk of 37% in their cohort while Roberts et al estimated risk at 42% for men and 33% for women by age 80.^42, 43^ These recent reports along with our findings suggest that some germline *CDH1* variant require other oncogenic molecular and/or environmental factors to drive DGC formation. This represents an opportunity for precision treatment strategies as we hypothesize that different variants may produce varied biological effects and targets for therapy. Finally, while five of our seven patients would have undergone genetic testing based on IGCLC recommendations to test DGC patients diagnosed before age 50, two did not meet criteria. A recent study by Lowstuter et al found that 65% of *CDH1* mutation carriers did not meet IGCLC guidelines for testing.^44^ This suggests that revisions will be necessary to improve the sensitivity of guidelines for genetic testing to identify germline *CDH1* carriers.

Previous analyses of early-onset gastric cancer have identified DGC as being associated with young age.^12, 45^ The high rate of DGC in Hispanic/Latino patients is consistent with the younger age of diagnosis in this cohort. However, the molecular mechanism behind early-onset carcinogenesis in this subgroup is unknown. As discussed above, the high rate of germline *CDH1* mutations in the Hispanic/Latino cohort may play a role. Previous studies in non-Hispanic/Latino cohorts showed that germline *CDH1* mutations occurs in about 1-3% of non-familial gastric cancer patients.^27–30^ In addition, the TCGA reported only two *CDH1* variants in 295 patients, and these were non-pathogenic.^12^ Other factors unrelated to Lauren classification and GS subtype clearly affect the etiology of early-onset gastric cancer since three of four very young (<35 years old) patients in our cohort were in the CIN group, with two of them having intestinal-type cancers. This will be an important area for future study, as the incidence of gastric cancer is rising in the United States only amongst young patients and thus will likely disproportionately affect the Hispanic/Latino population and exacerbate gastric cancer outcome disparities.^46^

While molecular classification systems proposed by the TCGA and others have provided new paradigms to study gastric cancers, the practical implications of this scheme for patient care remain elusive. Recently, Sohn et al. reported that the TCGA classification may provide both prognostic and predictive value in Korean patients.^47^ They found that EBV tumors had the best outcomes and GS cancers had the worst. Recently, Kim et al showed that immunotherapy was effective mainly in EBV or MSI-type tumors, while CIN and GS cancers were generally resistant.^48^ These findings have significant implications for the Hispanic/Latino cohort that we analyzed since 95% of the patients had either CIN or GS tumors. Using consensus clustering of RNA-seq data, we identified transcriptomic signatures that were prognostic and thus can aid in risk-stratification and treatment planning, as clinicians and patients contemplate the use and sequence of systemic therapies versus resection. Intriguingly, patients in the low-risk, favorable prognosis group had a gene signature suggestive of an activated immune response. Whether these patients will benefit from immune therapy is a possibility that should be tested.

Future studies of Hispanic/Latino populations will likely benefit from more refined definitions of ancestry mix. Hispanic and Latino groups encompass a geographically diverse population exhibiting significant genomic heterogeneity, due to the differential admixture of European, Indigenous American, and African populations. Previous studies have shown that ancestry proportions in Hispanic/Latino patients are associated with breast cancer incidence and outcomes.^49, 50^ However, while our study cohort is derived from patients living in North Texas, the country of origin is heterogeneous, as suggested by the relatively broad cluster seen in the ancestry analysis.

In conclusion, while gastric cancer outcome disparities may result from a combination of environmental exposures and socioeconomic factors, inherent tumor biology is also an essential component. Our study analyzing a large cohort of Hispanic/Latino gastric cancer patients is an important step in addressing the outcome disparity that these patients face by providing a genomic context for their disease. We have found that gastric cancers arising in Hispanic/Latino patients exhibit significantly different genomic landscapes than those developing in Asian and White patients. There is an enrichment for GS tumors and a high rate of germline *CDH1* mutations. Our findings should be considered in establishing guidelines for screening, genetic counseling, and treatment of Hispanic/Latino gastric cancer patients.

## Acknowledgements

S.C.W. is a UT Southwestern Disease-Oriented Scholar and an American College of Surgeons Research Faculty Fellow and is supported by a Society for Surgery of the Alimentary Tract Health Care Disparity Research Award and by the NIH/NCI (K08 CA222611). M.R.P. is a Dedman Family Scholar in clinical care. I.N. was supported by the National Center for Advancing Translational Sciences of the National Institutes of Health, under Award No. UL1TR001105. We acknowledge the assistance of the University of Texas Southwestern Tissue Resource, a shared resource at the Simmons Comprehensive Cancer Center, which is supported in part by the National Cancer Institute, under award number 5P30CA142543. The authors have no financial disclosures or conflicts of interest. We thank Drs. Herbert J. Zeh, III and Todd A. Aguilera for critical reading of the manuscript.

## Supplementary Figure Legend

**Supplementary Figure 1.**
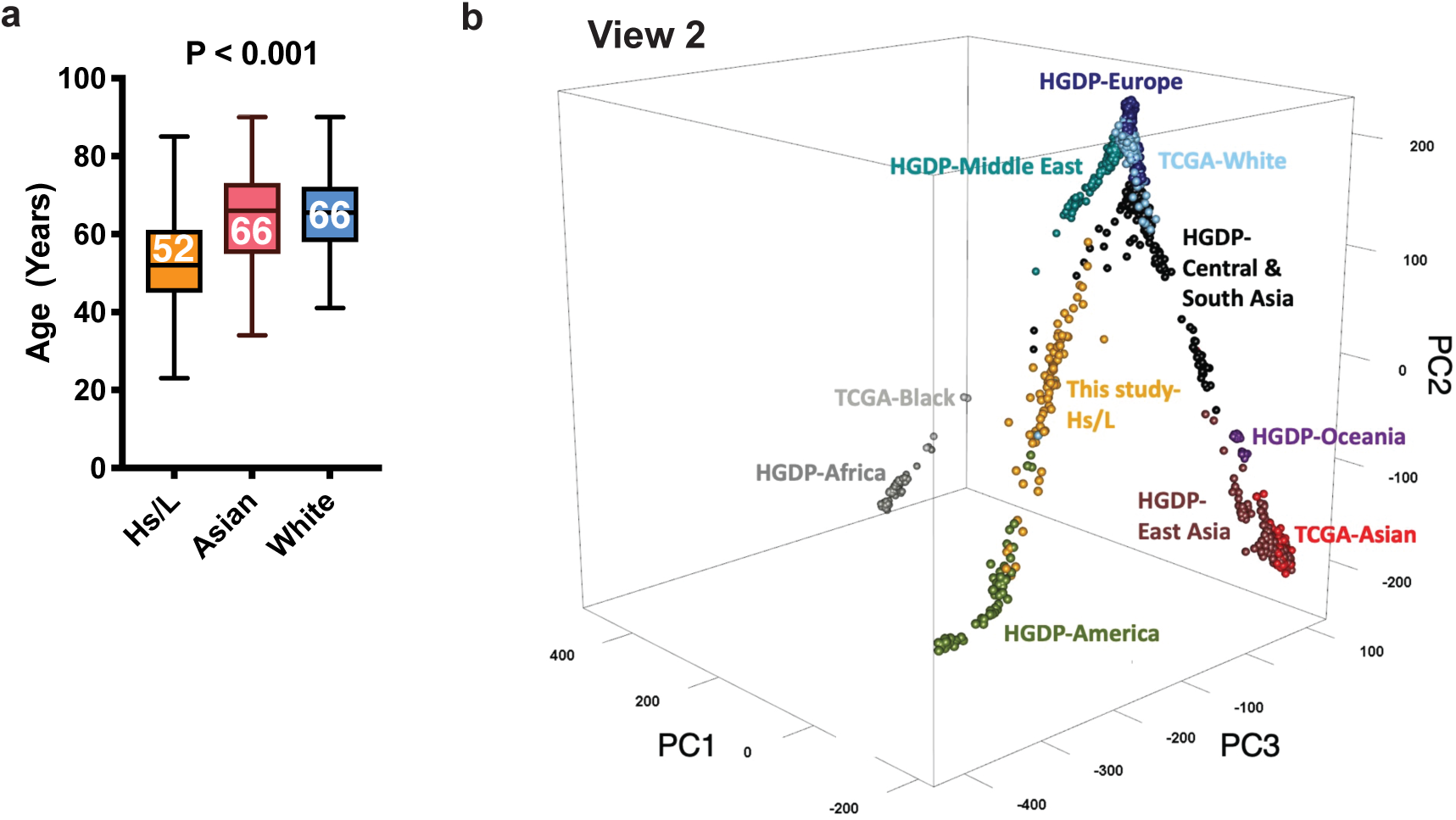
a. Age at the time of diagnosis of Hispanic/Latino (Hs/L) patients from this study, and of Asian and White patients analyzed by The Cancer Genome Atlas (TCGA). Horizontal lines, medians; boxes, interquartile ranges; whiskers, maximum and minimum values. P < 0.001. b. Second view of the principal component analysis of whole-exome sequencing data, as analyzed by Locating Ancestry from SEquence Reads to define patient ancestry of Asian and White patients analyzed by the TCGA and Hispanic/Latino patients from this study, as compared to reference from the Human Genome Diversity Project (HGDP).

**Supplementary Figure 2.**
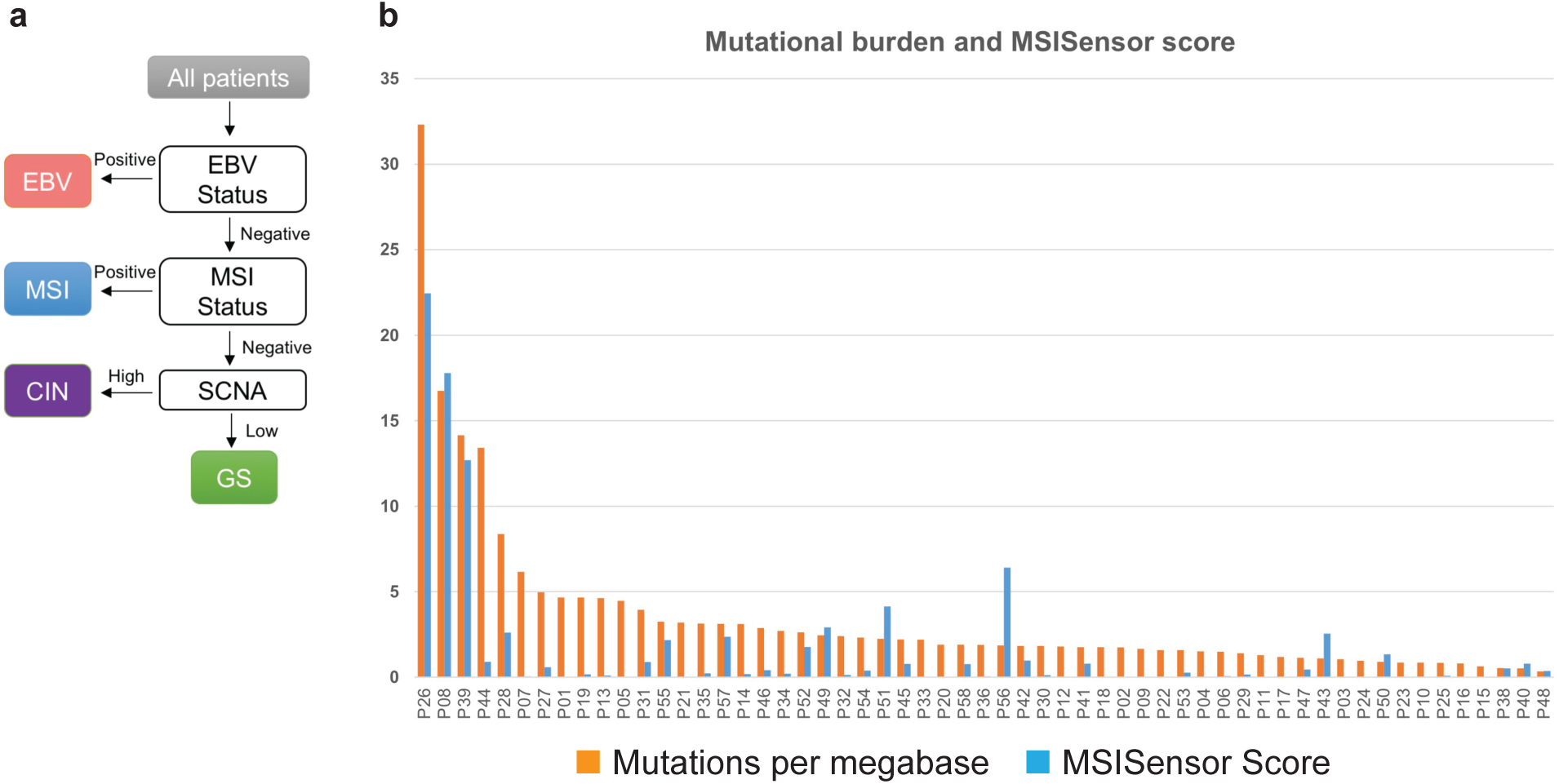
a. The Cancer Genome Atlas algorithm to categorize gastric cancer into four molecular subtypes: Epstein-Barr virus infected (EBV, red), microsatellite instability (MSI, blue), chromosomal instability (CIN, purple), and genomically stable (GS, green). SCNA = somatic copy number alterations. b. MSIsensor score. Whole-exome sequence data from each Hispanic/Latino cancer sample were analyzed with MSIsenor. Orange bar denotes total mutation burden per megabase. Blue bar denotes calculated MSIsensor score. Samples with score ≥ 10 were considered to be microsatellite unstable.

**Supplementary Figure 3.**
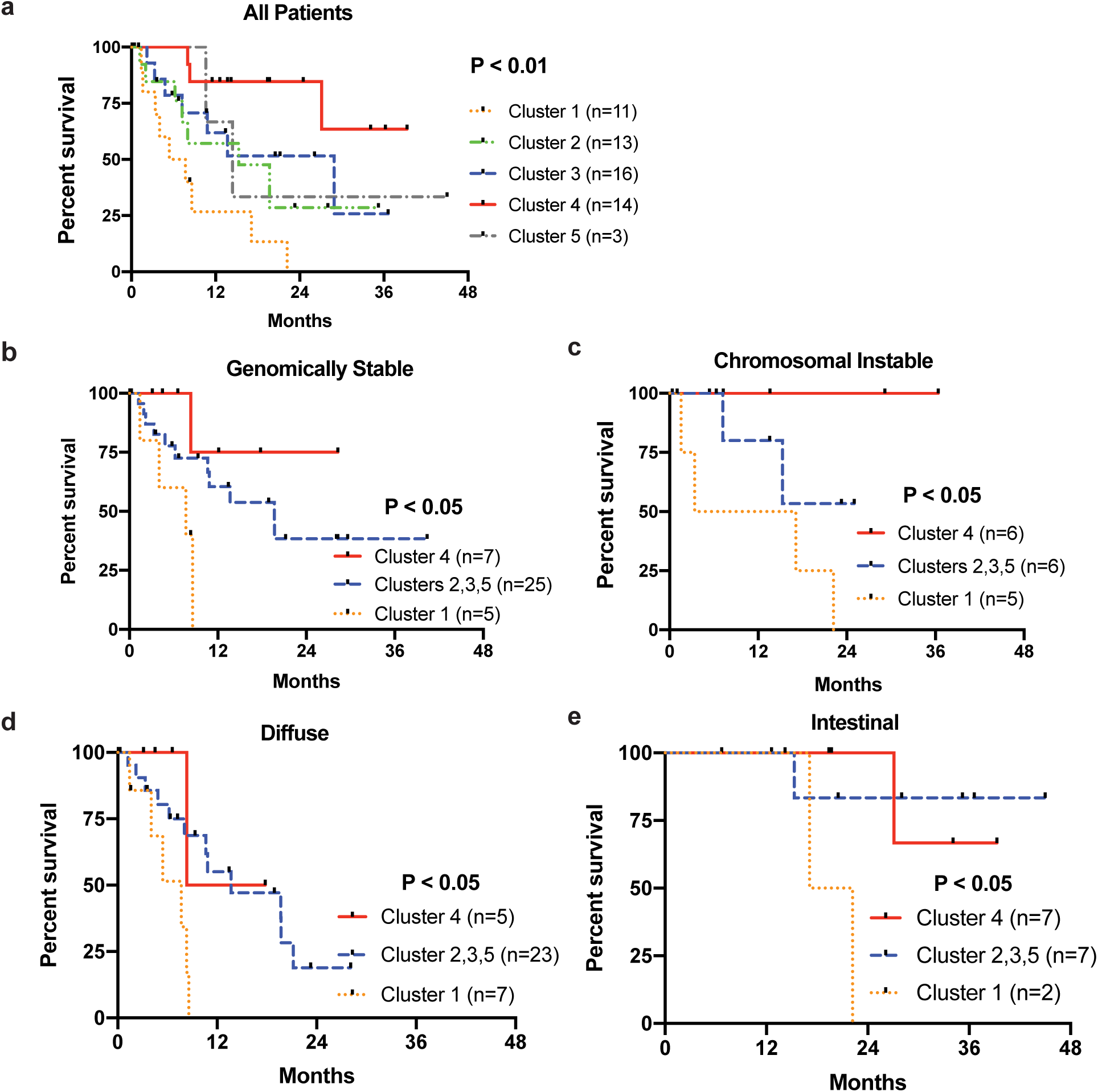
Subgroup analysis of overall survival for the Hispanic/Latino cohort. Kaplan-Meier curves comparing: a. all patients by individual mRNA clusters, P < 0.01, b. patients with genomically stable tumors, P < 0.05, c. patients with chromosomal instable tumors, P < 0.05, d. patients with diffuse-type tumors, P < 0.05, e. patients with intestinal-type tumors, P < 0.05.

**Supplementary Figure 4.**
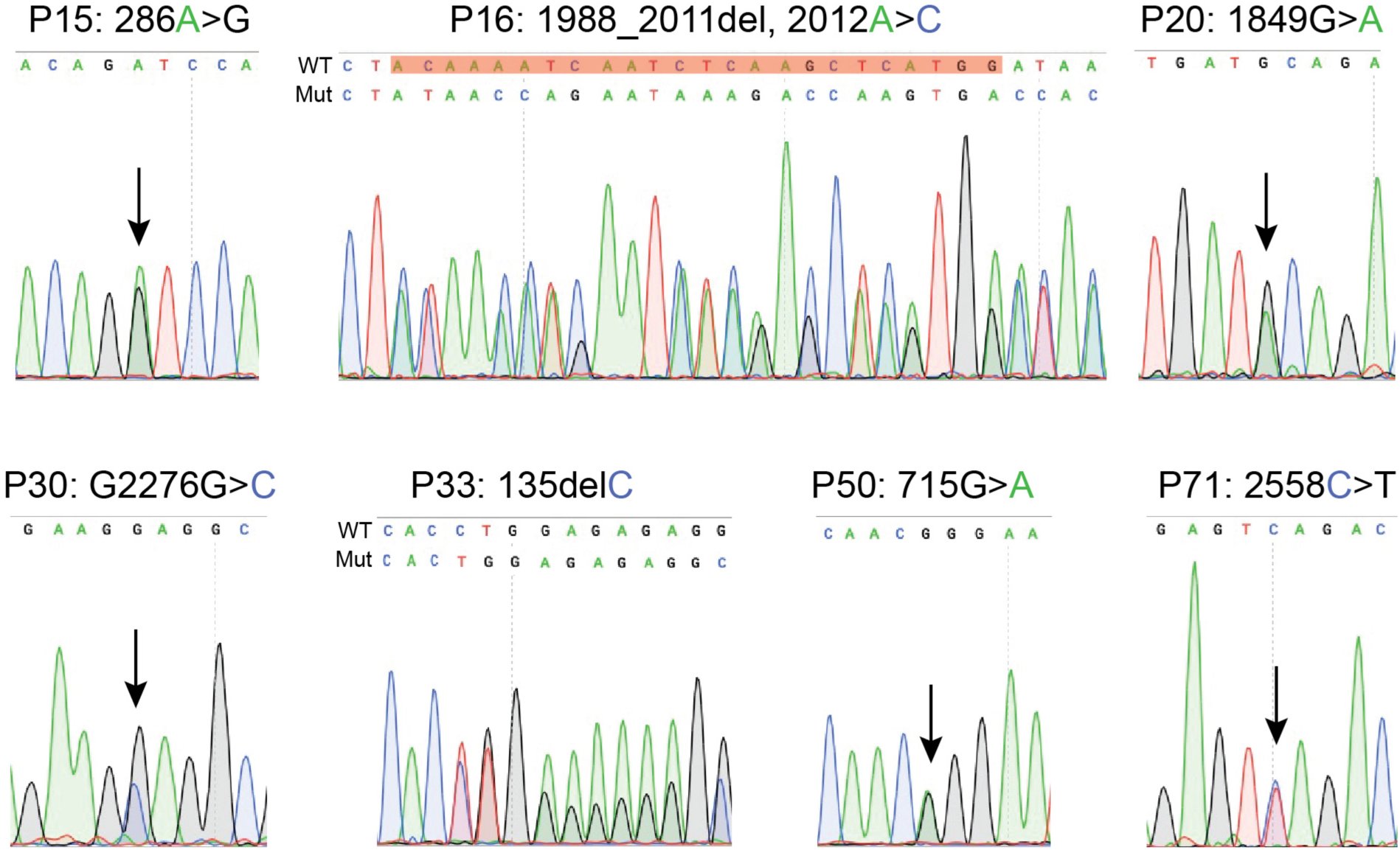
Chromatograms confirming germline *CDH1* mutations identified on whole-exome sequencing.

**Supplementary Figure 5.**
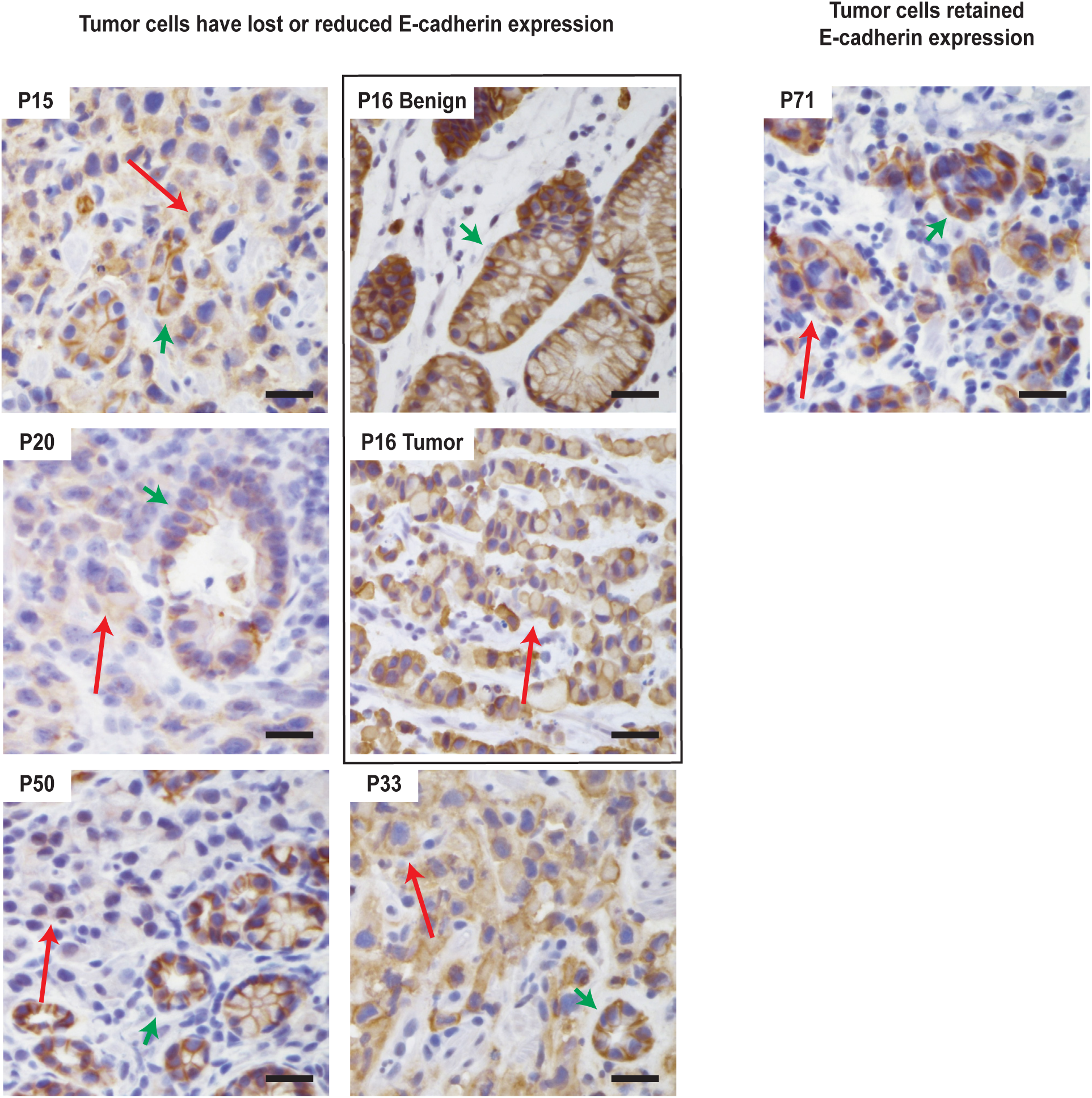
Immunohistochemistry for E-cadherin in the seven patients who have germline *CDH1* mutations. Red long arrows denote cancer cells, short green arrows denote normal stomach glands. P: patient. For P16, normal stomach and cancer cells are shown in two separate panels. Scale bar = 50 μm.

**Supplementary Figure 6.**
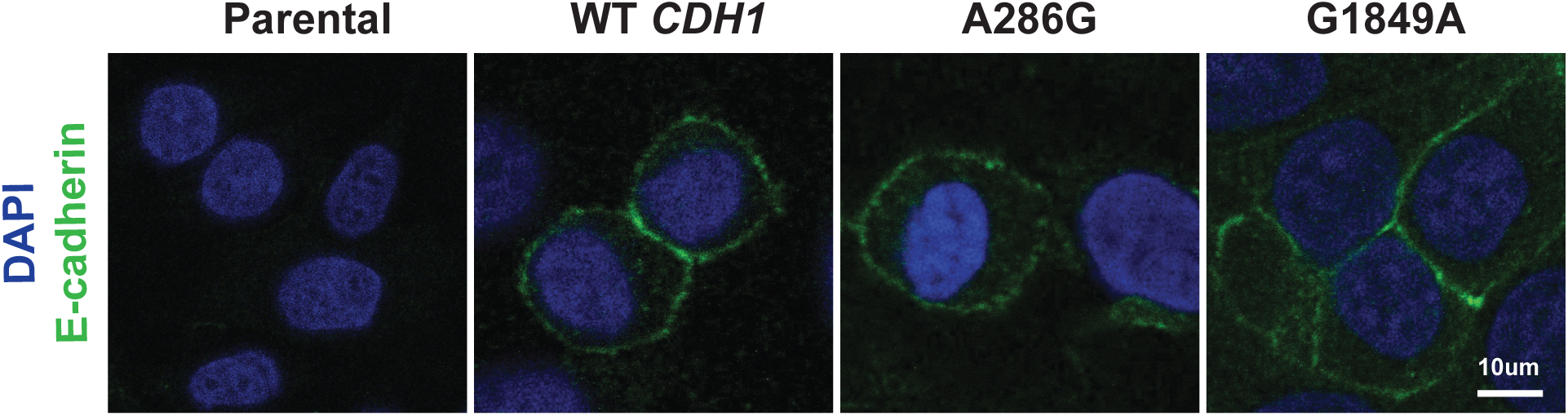
Immunofluorescence staining for E-cadherin in Chinese hamster ovary cells overexpressing wild-type (WT) *CDH1*, A286G, or G1849A variants, all of which have wild-type membranous localization. Scale bar = 10 μm.

## Supplemental Methods

### Whole-exome sequencing analysis

Libraries were made with the SureSelect XT Human All Exon V5+UTR kits (Agilent) and sequenced on HiSeqX (Illumina). We used FastQC v0.11.5_1_ and multiQC v1.2_2_ to check the quality of the raw FASTQ files and cutadapter v1.5 to remove adapter sequences._3_ The bwa-mem v0.7.15_4_ with default parameters was used to align to the hg19 reference genome._5_ PCR-duplicated reads were removed using Picard tools v2.9.0,_6_ then alignments were recalibrated using GenomeAnalysisToolkit (GATK) v3.7_7_ with known variant databases._8–10_

We used MuTect v1.1.7_11_ and Strelka2 v2.8.2_12_ to detect somatic variants for tumor and matched normal samples. For MuTect, alignment files were realigned using GATK, then were inputted into MuTect with the dbSNP_8_ and COSMIC databases._13_ Strelka2 was executed in somatic configuration with default parameters. We selected somatic variants that were detected by either MuTect or Strelka2 and annotated by ANNOVAR._14_ We further filtered variants that were not present within the exome capture kit. The variants present in the intronic or intergenic regions were also excluded. In addition, the variants with low variant allele frequency (VAF) with low quality supporting reads were filtered out. Specifically, variants with VAF ≤ 0.2 were excluded if they had less than 3 high quality supporting reads. We defined high quality supporting read as a read containing the mutated nucleotide with a base quality score of 30 or higher at the mutation as well as alignment quality score of 50 or higher given by the aligner for the entire read.

We used MutsigCV 1.4.1_15_ to identify statistically significant recurrent somatic variations. Full exome coverage data from MutsigCV website was used for this test. We used *p* value ≤ 0.05 as a cutoff for determining the significance of the recurrent mutations. We used GenVisR to generate a waterfall plot to visualize a pattern of recurrent variations in our cohort._16_

Germline variations were identified with GATK Haplotypecaller and Strelka2. For GATK Haplotypecaller, we used recommended parameters and a variant recalibration stage with known variant databases._8–10, 17_ Because we focused on few specific cancer related genes, we used the union of the two germline mutation profiles to increase the sensitivity, then we manually inspected potential germline variations using IGV._18_ We also annotated the germline mutations using ANNOVAR, then selected a subset of mutations as potentially pathogenic by the following two criteria: 1) predicted as deleterious by SIFT_19_ and PolyPhen2_20_ and 2) having a lower population frequency than 0.01 in gnomAD (https://gnomad.broadinstitute.org).

### RNA sequencing and analysis

Libraries were made with TruSeq RNA Access Library kit (Illumina) and sequenced on HiSeq2500 (Illumina). We used FastQC, multiQC, and cutadapt for quality control and preprocessing. STAR v2.4.2a_21_ was used to align to a GRCh38_P01 reference with Genecode v22 gene annotation, then HTSeq 0.6.0_22_ was used to generate counts for gene expression quantification. The same parameters used in the TCGA STAD project_23_ were used. DESeq2_24_ was used to detect differentially expressed genes (DEG).

Consensus clustering using gene expression data was performed with ConsensusClusterPlus._25_ The raw read counts were normalized using voom function in the limma package. Various k (number of clusters) from 2 to 20 were tested and k = 5 was selected based on Cophenetic correlation coefficient. We performed DEG analysis between the identified cluster 1 and clusters 2+3+4+5 and cluster 4 and clusters 1+2+3+5 using the DESeq2 and used GSEA for gene set enrichment analysis._26_

FusionCatcher_27_ and STAR-Fusion_28_ were used with default parameters to detect gene fusion events. Fusion events identified by both algorithms were used for further analysis. Non-clipped raw FASTQ data and Ensembl v89 database_29_ were used as input and database, respectively. We visualized and manually inspected fused transcripts using supporting reads provided by the FusionCatcher and UCSC genome browser._30_

### Ancestry analysis

To identify each sample’s inherited genetic characteristics, we performed an ancestry inference using Locating Ancestry from SEquence Reads (LASER) to analyze whole-exome sequencing data_31_ with default parameters. LASER constructs a reference principal component (PC) space with a set of reference individuals and places test samples into the PC space. Ancestry of each sample can be inferred using distances in the PC space between the sample and the reference individuals. We downloaded and used a reference PC space data from the LASER website. The reference PC space was constructed with Human Genome Diversity Project_32_ data that contained 938 reference individuals from various ethnic groups. Then we calculated the first 4 PCs for each normal Hispanic/Latino sample and mapped it to the PC space.

### Metagenomics for Epstein-Barr virus and Helicobacter pylori

We used PathoScope 2.0_33_ with parameters (-b very-sensitive-local -m hi -k 100 -t 50 -L 101 -s 0.95 --adjreflen --reuse) to identify EBV infections using whole-exome and RNA sequencing data. We used the target microbial database (PathoDB) available from PathoScope 2.0 release, which was built from NCBI nr (non-redundant) nucleotide database as of 2014._34_ To increase sensitivity, we performed the metagenomics analysis on both WES data and RNA-seq data.

### Determining microsatellite instability

We used MSISensor_35_ with default parameters to predict the MSI status by calculating and comparing length distributions of microsatellites between tumor and normal sample. The MSISensor calculated a score for each sample to determine the MSI status (e.g., higher scores indicate the sample is more likely to have MSI). If we have both of normal and blood samples for a patient, we did two tests and averaged the scores. Then we chose a cutoff 10 based on a pan-cancer MSI assessment study using the MSISensor._36_

### Determining somatic copy number alterations

We used CNVkit_37_ to perform copy number alteration (CNA) analysis for whole-exome sequencing data. Because our cohort was sequenced by two different vendors (DNA Link, Inc (San Diego) and Admera Health (New Jersey)) we divided the cohort into two batches based on the vendors and ran the CNVkit separately. For each batch, a pooled reference panel was built using normal samples, then somatic CNAs were called for each tumor sample. The CNA calling results were merged then GISTIC2_38_ was used to identify recurrent CNA regions. CNA regions with |CNA value| < 0.1 were filtered out. We adopted a method from Ichikawa et al._39_ and (number of CNA regions > 41) was defined as a cutoff to stratify between genomically stable (GS) and chromosomal instability (CIN) subtypes.

## Supplementary Tables

**Supplementary Table 1:** Clinicopathologic information for each patient sequenced for this study.

**Supplementary Table 2:** Details of *CLDN18-ARHGAP* fusion events.

**Supplementary Table 3:** Clinical and molecular information for each mRNA cluster.

**Supplementary Table 4:** Published reports with rates of *CDH1* germline mutations in non-familial gastric cancer patients.

